# Reassortant strains of infectious bursal disease virus (IBDV) belonging to genogroup A3B1 predominate in British broiler chicken flocks

**DOI:** 10.1101/2024.04.24.590994

**Authors:** Vishwanatha R. A. P. Reddy, Carlo Bianco, Christopher Poulos, Andrew J. Brodrick, Salik Nazki, Alex Schock, Andrew J. Broadbent

**Affiliations:** The Pirbright Institute, Ash Road, Woking, GU24 0NF, UK; School of Life Sciences, Keele University, Keele, ST5 5BG, UK; Animal and Plant Health Agency (APHA) Lasswade, Penicuik, Midlothian, EH26 0PZ, UK; School of Veterinary Medicine and Science, University of Nottingham Campus, Sutton Bonington, LE12 5RA, UK; Department of Animal and Avian Sciences, University of Maryland, College Park, MD, 20742, USA; Pandemic Sciences Institute, Nuffield Department of Medicine, University of Oxford, Oxford, UK; Chinese Academy of Medical Sciences Oxford Institute, Nuffield Department of Medicine, University of Oxford, Oxford, United Kingdom

**Keywords:** infectious bursal disease virus, IBDV, diversity, phylogenetics

## Abstract

As part of ongoing epidemiological surveillance for infectious bursal disease virus (IBDV), the hypervariable region (HVR) of the VP2 capsid gene encoded by segment A, and a region of the VP1 polymerase gene, encoded by segment B, were sequenced from 20 IBDV-positive bursal samples obtained in 2020 and 2021, from 16 commercial British broiler farms. Of the 16 farms, none contained very virulent (vv) strains belonging to genogroup A3B2, but 5/16 (31%) contained strains of genogroup A3B1, demonstrating birds were infected with reassortant strains containing a vv segment A and a non-vv segment B. In addition, 3/16 (19%) farms contained vaccine or classical strains belonging to genogroup A1B1, and 8/16 (50%) were co-infected with both genogroup A1B1 and A3B1 strains. Therefore, a total of 13/16 (81%) of the farms contained genogroup A3B1 reassortant viruses, the majority of which 8/13 (62%)) were found to be co-infected with genogroup A1B1 strains. Moreover, of the flocks containing reassortant strains, 5/13 (38%) had HVR mutations Q219L, G254D, D279N, and N280T, consistent with a recently described Western European clade, but 8/13 had other mutations or no mutations, demonstrating that multiple clades were present in the samples. Taken together, vv strains were not detected in the British broiler flocks we sampled, whereas reassortant strains predominated, which belonged to different clades, and were frequently found in samples that were also infected with genogroup A1B1 strains.

## Introduction

Infectious bursal disease virus (IBDV), a member of the *Avibirnavirus* genus in the *Birnaviridae* family, is a highly contagious and immunosuppressive virus that infects poultry worldwide, and is ranked among top infectious problems of chickens (1). The virus is non-enveloped, with a genome comprised of double stranded RNA that is divided into two segments, segment A (3.2 Kb) and segment B (2.8 Kb), enclosed within a capsid with icosahedral symmetry (1). Segment A has two partially overlapped open reading frames (ORFs). ORF A1 encodes the non-structural viral protein VP5 that is reported to be involved in virus egress (2), and ORF A2 encodes a large polyprotein that undergoes cleavage by the protease VP4 to yield VP2, VP4, and VP3 (3). VP2 is the capsid protein, and VP3 is a multifunctional protein that binds the dsRNA genome and may help form a scaffold between the genome and the capsid (4, 5). Segment B has one ORF that encodes the RNA dependent RNA polymerase (VP1) enzyme, which replicates the viral genome (6).

The population of IBDV field strains has traditionally been divided into different pathotypes: classical (c-) field strains, antigenically variant field strains, and very virulent (vv) field strains (1), and both segments A and B contribute to the pathogenicity (7). The c-field strains were first identified in the 1960s and continue to circulate today. These strains are immunosuppressive, and can cause clinical signs, such as ruffled feathers, drooping wings, lethargy, soiled vent feathers or diarrhoea. Antigenically variant field strains first emerged in the 1980s in the USA, and while they may not cause severe clinical signs, they can cause immunosuppression in infected birds, exacerbating secondary infections and hindering the response to other vaccination programs. The vv strains also emerged in the 1980s and cause a severe acute disease with high mortality (1). In addition to these field strains, live vaccine strains may also be detected in flocks.

The VP2 capsid is known to be an important immunodominant protein of IBDV and is the major target of neutralizing antibodies. Within VP2, there is a so-called “hypervariable region” (HVR) located between amino acids 220 to 330, which is subject of the most intense immune selection pressure and antigenic drift. The global population of IBDV strains have been divided into different genogroups based on the sequence diversity of the HVR. Initially, 7 A genogroups were reported, then 8, and now a 9^th^ genogroup has recently been described (8–10). Classical strains belong to genogroup A1, antigenically variant strains, particularly those in the USA and China belong to genogroup A2, and very virulent strains belong to genogroup A3. Other strains isolated in South America belong to genogroup A4, recombinant strains identified in Mexico belong to genogroup A5, and some strains identified in Italy and Eastern Europe belong to genogroup A6. Strains isolated in Australia belong to genogroups 7 and 8 (8), whereas strains recently isolated in Portugal belong to genogroup A9 (9). Furthermore, within the HVR, there are four hydrophilic loops of amino acids in the tip of the VP2 molecule, termed P_BC_, P_DE_, P_FG_, and P_HI._ These loops are reported to contribute to IBDV virulence and determine the antibody neutralization profile of the virus (11–15). Continued antigenic drift at these sites is driving ongoing evolution of the genogroups. Epidemiological studies typically also target a region of the VP1 gene to further classify IBDV. VP1 is encoded by segment B, and the population of IBDV strains have been divided into 5 genogroups based on this segment (8). IBDV strains are therefore currently typed based on their A and B genogroups, for example classical strains and many vaccine strains belong to genogroup A1B1, US variant strains are A2B1, and vv strains are A3B2. Owing to the segmented nature of the genome, it is possible for IBDV to undergo reassortment in co-infected birds, where the offspring virus may contain a mixture of A and B genome segments from both parental strains. It is therefore important in molecular epidemiology studies to sequence both segments, and numerous detections of reassortant strains have been reported across the globe, between strains belonging to different genogroups, and even between serotypes. In 2020 and 2023, reassortant IBDV strains with a vv segment A and non-vv segment B belonging to genogroup A3B1 were found to be widespread in chicken flocks in Europe. Moreover, some of the viruses had undergone antigenic drift and accumulated HVR mutations, to form a Western European clade of reassortant strains (9, 16–20).

To further study the landscape of IBDV in the Great Britain, and establish the status of endemic, evolving/emerging, and exotic variants, we conducted an active epidemiological transdisciplinary investigation. To this end, we obtained 20 bursal samples in 2020 and 2021, from birds belonging to 16 commercial British broiler farms, which showed histological lesions consistent with an IBDV infection. We sequenced the VP2 HVR and a region of the VP1gene and conducted phylogenetic analysis and alignments.

## Materials and Methods

### Sampling protocol

Bursal samples were collected between 2020 and 2021 from British broiler farms through the endemic disease surveillance network of the Animal and Plant Health Agency (APHA) which is based on voluntary contribution from private veterinarians. Samples were collected by private veterinary surgeons during routine on-farm post-mortem examinations because of a suspicion of an IBDV infection. Samples from each farm were assigned a number to anonymise the results. The age of the birds and the type of vaccination used were also collected. To maintain confidentiality, no additional data were gathered. A representative number of bursae was fixed in formalin and processed for routine histopathological investigations at the APHA, Lasswade, and a morphological diagnosis was established. The remainder of the tissue was pooled and frozen at -80°C and shipped to the Pirbright Institute on dry ice, whereupon it was stored at -80°C until processing in the laboratory. Where multiple samples were obtained from the same farm, each sample was also assigned a letter. In case of farm 3, the three samples (3a, b, and c) represented three individual bursae from the same house, and each was sequenced separately. In case of farms 4 and 8, the two samples (4a and b, and 8a and b) each represented a pool of three bursae obtained from two different houses on the same farm at the same point of production, but only one bursal sample from each pool was sequenced.

### RNA extraction, partial, full-length segment amplification, and sequencing

Total RNA was extracted from 30 mg of one bursa, using an RNeasy kit (Qiagen) according to the manufacturer’s instructions. Complementary DNA (cDNA) was generated using SuperScript III Reverse Transcriptase (Invitrogen) and a random primer. The reaction constituents and conditions were consistent with the manufacturer’s instructions. Previously reported primers,743-F (5’-GCCCAGAGTCTACACCAT -3’) and 1331-R (5’-ATGGCTCCTGGGTCAAATCG-3’) were used to amplify a 579-bp fragment of the HVR of the VP2 of segment A (10), and B-168A-F (5’-CATAAAGCCTACAGCTGGAC-3’) and B-889-R (5’-GTCCACTTGATGACTTGAGG-3’) were used to amplify a 722bp region of segment B in a polymerase chain reaction (PCR) (21). Amplification of both VP2 and VP1 was performed using same reaction conditions described previously (10). Briefly, following an initial denaturation at 94°C for 2 minutes, DNA was amplified in 35 cycles comprised of denaturation at 95°C for 30 seconds, annealing at 57°C for 1.5 minutes, and extension at 72°C for 1.5 minutes with a final extension at 72°C for 5 minutes. PCR products were subjected to electrophoresis in tris-borate EDTA (TBE) gel containing 2% agarose, bands were excised and purified using the Illustra GFX kit (GE) according to manufacturer’s instructions. Both VP2 and VP1 sequences strands were sequenced by GATC BiotechAG (Konstanz, Germany) by Sanger sequencing.

### Phylogenetic analysis

To perform phylogenetic analysis, the HVR and partial VP1 sequences were aligned with VP1 and VP2 sequences from representative IBDV strains (8, 10). Alignments were performed with the ClustalW algorithm in MEGA software version 7 (22). Phylogenetic trees were constructed using the Neighbour joining method in MEGA software version 7, with 1,000 bootstrap replications and a Kimura 2 (K2) + G parameter model Bootstrap values lower than 75% were considered as non-significant. The generated phylogenetic data were visualised using the Interactive Tree of Life Tool (iTOL) (23).

### Structural modelling

The structure of the HVR monomer from a Western European clade reassortant virus was modelled using a modified version of AlphaFold v2.3.2. The sequence from which the structure was predicted was generated by adding mutations Q219L, G254D, D279N, and N280T to the UK661 VP2 sequence *in silico*. The resultant prediction was then downloaded and processed using PyMol (v2.5; Schrödinger) to visualize the structure and highlight mutation sites. To view the mutation sites in the context of the VP2 trimer, 3 copies of the UK661 VP2 sequence (NCBI reference sequence NC004178) were processed with the AlphaFold multimer model, with an allowance of 20 recycles. The prediction was then aligned to a published VP2 trimer extracted from an IBDV capsid structure solved by Cryo-EM (24), obtained from the Research Collaboratory for Structural Bioinformatics (RCSB) Protein Data Bank (accession code 7VRN), to determine how reasonable the model was. The alignment showed agreement between the structures (root mean square deviation (RMSD) = 1.27Å). The predicted trimer was then processed with PyMol to highlight the position of the mutations of the Western European clade reassortant virus.

### TA cloning to separate mixed infections

The cDNA from the 20 bursal samples was subject to PCR to amplify the HVRs, and partial VP1 sequences as described. The Taq DNA polymerase added a single deoxyadenosine to the 3’ end of the products, which were then ligated into a vector containing a complimentary thymine provided in a TA cloning kit (ThermoFisher Scientific). Competent DH5α E. coli cells (New England Biosciences) were transformed and streaked onto agar plates with antibiotic selection, where the bacteria formed colonies. If the PCR products contained a mixture of sequences, each were separated into a different colony. 1-10 colonies were picked for each amplified HVR product, and each VP1 product, for each of the 20 field samples, and subject to colony PCR using a PlateSeq Kit (Eurofins Genomics).

## Results

### Twenty bursal samples were obtained from 16 British commercial broiler farms between 2020 and 2021

All the flocks received live attenuated vaccines (LAVs) belonging to genogroup A1B1, and one flock received an HVT-vectored vaccine followed by a LAV. Birds were vaccinated with LAV at 17-21 days of age, and humanely euthanised at 25-55 days of age for postmortem analysis. Histopathological data was unavailable for one sample. Out of the remaining 19 samples, nine showed bursal atrophy, two showed non-specific medullary lymphoid depletion, three had bursitis, and five showed both bursitis and bursal atrophy (Table 1). From three farms (3, 4 and 8), multiple samples were received from the same flocks.

**Table 1.** Details on the 20 bursal samples obtained from 16 British broiler farms.

| **Farm number<sup>+</sup>** | **Vaccination age (days)** | **IBDV Vaccine** | **Age at necropsy (days)** | **Pathology** |
| --- | --- | --- | --- | --- |
| 1 | 17 | LAV<sup>#</sup> 1 | 30 | Bursal atrophy |
| 2 | 17 | LAV 2 | 34 | Bursitis / Bursal atrophy |
| 3 (3 samples 3a, 3b and 3c) | 18 | LAV 3 | 30 | Bursitis |
| 4 (2 samples 4a and 4b) | 17 | LAV 2 | 37 | Bursitis / Bursal atrophy |
| 5 | 17-18 | LAV 4 | 31 | Bursal atrophy |
| 6 | 18 | LAV 2 | 55 | Bursal atrophy |
| 7 | 18 | LAV 2 | 25 | Medullary lymphoid depletion |
| 8 (2 samples; 8a and 8b) | 21 | LAV 5 | 38 | Bursitis / Bursal atrophy |
| 9 | 18 | HVT vectored vaccine + LAV 2 | 43 | Bursal atrophy |
| 10 | 22 | LAV 6 | 27 | Medullary lymphoid depletion |
| 11 | 17 | LAV 2 | 35 | Bursal atrophy |
| 12 | 17 | LAV 2 | 35 | Bursal atrophy |
| 13 | ND* | LAV 3 | 34 | Not done |
| 14 | 17 | LAV 2 | 37 | Bursal atrophy |
| 15 | 17 | LAV 2 | 32 | Bursal atrophy |
| 16 | 17 | LAV 2 | 33 | Bursal atrophy |

\*Not Disclosed (ND)
<sup>#</sup>Live Attenuated Vaccine (LAV); a total of 6 LAVs were used by the farms in this study, all belonged to genogroup A1B1
<sup>+</sup> Each set of samples consisted of three bursae from three different birds thought to be infected with IBDV

### Reassortant IBDV viruses were identified belonging to genogroup A3B1

Consensus sequences were obtained of the HVR of the VP2 gene encoded by segment A, and a region of the VP1 gene encoded by segment B from twenty samples, and were used to construct a phylogenetic tree for both segments A and B. According to the phylogenetic analysis based on the sequence of the VP2 HVR, 12/20 sequences (60%) belonged to genogroup A3 (vv strains) (samples 1, 3b, 3c, 4a, 4b, 5, 7, 9, 10, 12, 13 and 14), and 8 sequences (40%) belonged to genogroup A1 (classical or vaccine strains) (samples 2, 3a, 6, 8a, 8b, 11, 15 and 16) (Figure 1), whereas the phylogenetic analysis of the partial VP1 sequences revealed that all 20 sequences clustered with genogroup B1 (classical or vaccine strains) (Figure 2). Taken together, 12/20 contained reassortant strains belonging to genogroup A3B1, and 8/20 contained vaccine or classical strains belonging to genogroup A1B1. At the farm level, based on the phylogenetic analysis, of the 16 farms, 9/16 samples contained reassortant strains belonging to genogroup A3B1, 6/16 samples contained vaccine or classical strains belonging to genogroup A1B1, and one farm contained both A3B1 and A1B1 strains. No very virulent strains (genogroup A3B2) were detected.

**Figure 1.**
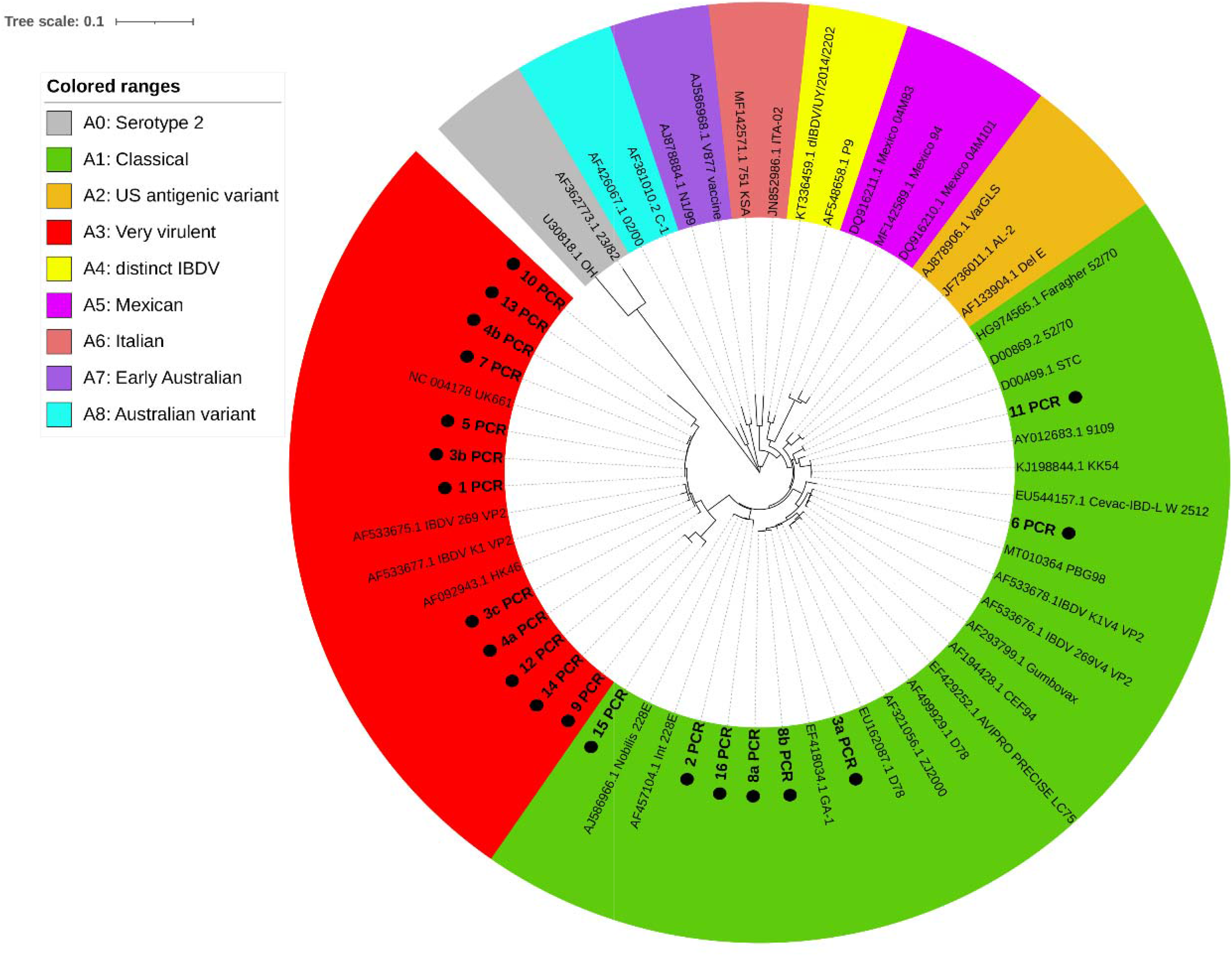
Phylogenetic tree based on the HVR of the VP2 gene, encoded by segment A. The tree was constructed using the consensus HVR nucleotide sequences obtained from Sanger sequencing of the PCR products from the 16 British broiler farms (labelled “1 PCR-16 PCR”), and the nucleotide sequences of reference IBDV strains in GenBank (indicated by the Accession numbers). The tree was constructed using the Neighbour-joining method in the MEGA software (version 7) with 1,000 bootstrap replications and a Kimura 2 (K2) + G parameter model. Bootstrap values lower than 75% were considered as non-significant. The phylogenetic data were visualised using the Interactive Tree of Life Tool (iTOL) (23), and the tree was divided into 9 sections, each depicted in a different colour, and representing genogroups A1-8 within serotype 1, as well as serotype 2 in grey, as indicated by the key. Each British field sample obtained is indicated by a black dot.

**Figure 2.**
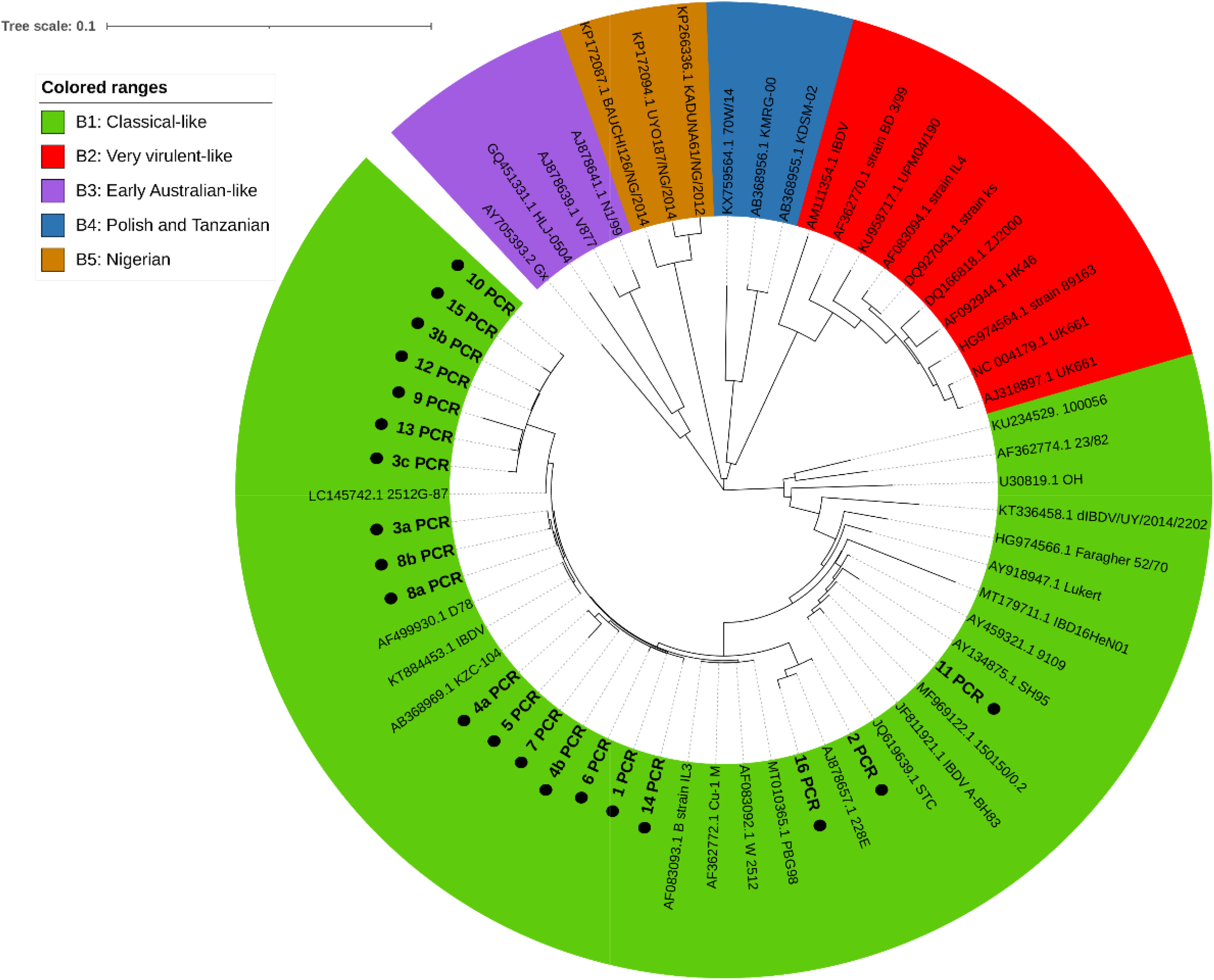
Phylogenetic tree based on a region of the VP1 gene, encoded by segment B. The tree was constructed using the consensus VP1 nucleotide sequences obtained from Sanger sequencing of the PCR products from the 16 British broiler farms (labelled “1 PCR-16 PCR”), and the nucleotide sequences of reference IBDV strains in GenBank (indicated by the Accession numbers). The tree was constructed using the Neighbour-joining method in the MEGA software (version 7) with 1,000 bootstrap replications and a Kimura 2 (K2) + G parameter model. Bootstrap values lower than 75% were considered as non-significant. The phylogenetic data were visualised using the Interactive Tree of Life Tool (iTOL) (23), and the tree was divided into 5 sections, each depicted in a different colour, and representing genogroups B1-5, as indicated by the key. Each British field sample obtained was indicated by a black dot.

### Reassortant strains co-circulated with classical/vaccine strains

While the phylogenetic trees were constructed based on the consensus sequences, further analysis of the chromatogram traces from the Sanger sequencing revealed that some showed a mixed trace, indicating that we had co-amplified a mixture of IBDV sequences from each of these bursal samples. To separate the traces, all 40 PCR products (20 of the HVR (Segment A) and 20 of the VP1 (Segment B)) were subject to TA cloning, and transformed into *E. coli,* which were then streaked onto agar plates to form colonies, each colony containing only one amplified product. For each cloned PCR product, 1-10 colonies were picked and sequenced, totalling 138 colonies for segment A and 112 colonies for segment B (Figure 3 and Supplementary Figures 1 and 2). Of the segment B sequences, all were found to be from genogroup B1 (vaccine or classical strains), with no genogroup B2 (vv) segments detected, even at a sub-consensus level. Of the segment A sequences, 10/ 20 (50%) had some colonies with sequences that corresponded to genogroup A3 and some that corresponded with genogroup A1, suggesting these birds were co-infected with reassortant viruses and classical or vaccine strains. The samples that had a mixture of A3 and A1 sequences included seven samples previously grouped as A3 based on the consensus sequences (samples 1, 3b, 3c, 4a, 4b, 5, and 10) and three samples previously grouped as A1 based on the consensus sequences (samples 6, 15, and 16). Taken together, our data suggest that 5/20 bursal samples sequenced contained genogroup A1B1 strains only (samples 2, 3a, 8a, 8b, and 11), 5/20 contained genogroup A3B1 reassortant strains only (samples 7, 9, 12, 13, and 14), and 10/20 were co-infected with both genogroup A1B1 and A3B1 strains (samples 1, 3b, 3c, 4a, 4b, 5, 6, 10, 15, and 16). At the farm level, 3/16 (19%) of the farms contained genogroup A1B1 strains, 5/16 (31%) contained genogroup A3B1 reassortant strains, and 8/16 (50%) were co-infected with both genogroup A1B1 and A3B1 strains. Therefore, a total of 13/16 (81%) of the farms contained reassortant viruses, the majority of which (8/13 (62%)) were found to be co-amplified with sequences of genogroup A1B1 strains.

**Figure 3.**
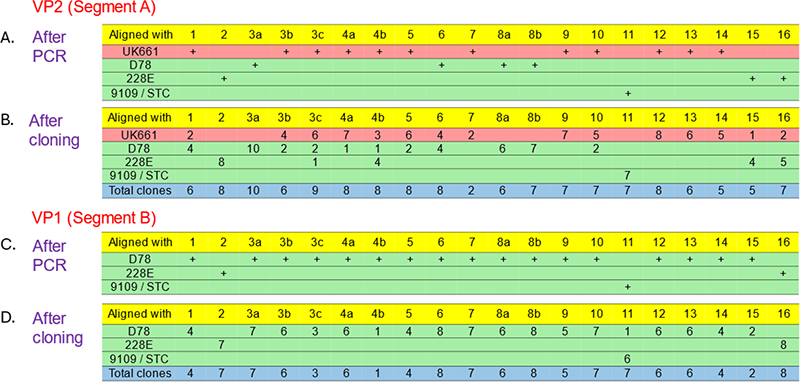
Separation of PCR products by TA cloning. The consensus nucleotide sequences of the VP2 HVR PCR products from the 20 bursal samples obtained from 16 British broiler farms (labelled 1-16) were grouped (+ symbols) based on their alignments with the vv strain UK661 (genogroup A3B2), depicted in red, and four strains of genogroup A1B1: classical vaccine strains D78 and 228E, and classical field strains 9109 and STC (depicted in green) (A). The PCR products were then subject to TA cloning, and transformed into *E. coli,* which were then streaked onto agar plates to form colonies. For each PCR product, 1-10 colonies were picked and sequenced (total number of colonies sequenced per sample depicted in blue). The number of colonies that contained sequences that grouped with either UK661, D78, 228E, or 9109/STC are shown (B). The consensus nucleotide sequences of the VP1 PCR products from each sample were similarly grouped (C), and subject to TA cloning, colony sequencing, and grouping (D).

### Antigenic drift mutations were identified in the British IBDV strains, consistent with the Western European branch of reassortant viruses

The sequences of segments A and B were translated *in silico* and the amino acid sequences aligned using ClustalW (MEGA software version 7) and compared to either the vv strain UK661 (Accession number NC004178), or the D78 classical vaccine strain (Accession number EU162087). Of the 12 consensus sequences of the A3B1 reassortant strains, three contained HVR antigenic drift mutations Q219L, G254D, D279N, and N280T (Figure 4). These mutations have previously been characterized in a Western European clade of reassortant viruses, demonstrating this branch is also present in Great Britain. This genetic signature was also present in two of the co-infected samples based on the colony sequencing (Supplementary Figure 1), meaning a total of 5 of the reassortant viruses were from this clade. At the flock level, 5/13 (38%) of the farms with reassortant viruses had strains from this clade. These mutations were located in the first three hydrophilic loops (P-BC, P-DE, and P-FG) (defined as per (11)) and were present both on the side of the HVR, and on the axial tip (Figure 5). In addition, 2/13 (15%) of the farms with reassortant viruses had strains containing T250S and S251I mutations in the P-DE loop compared to UK661, and the remaining 6/13 (46%) of the farms with reassortant viruses had either no mutations compared to UK661, or sporadic point mutations (Figure 4 and Supplementary Figure 1). No mutations were observed in the fourth hydrophilic loop (P-HI), in the consensus sequences of either viruses containing segment As from genogroup A3 (Figure 4), or genogroup A1 (Figure 6). The sequences of segment B were also aligned to UK661 (Accession number AJ318897.1), and 19/20 (95%) had mutations T145N, D146E, and N147G, and one (sample 11) had T145N, D146E, and N147D (Figure 7 and Supplementary Figure 2), indicating they were all from non-vv strains.

**Figure 4.**
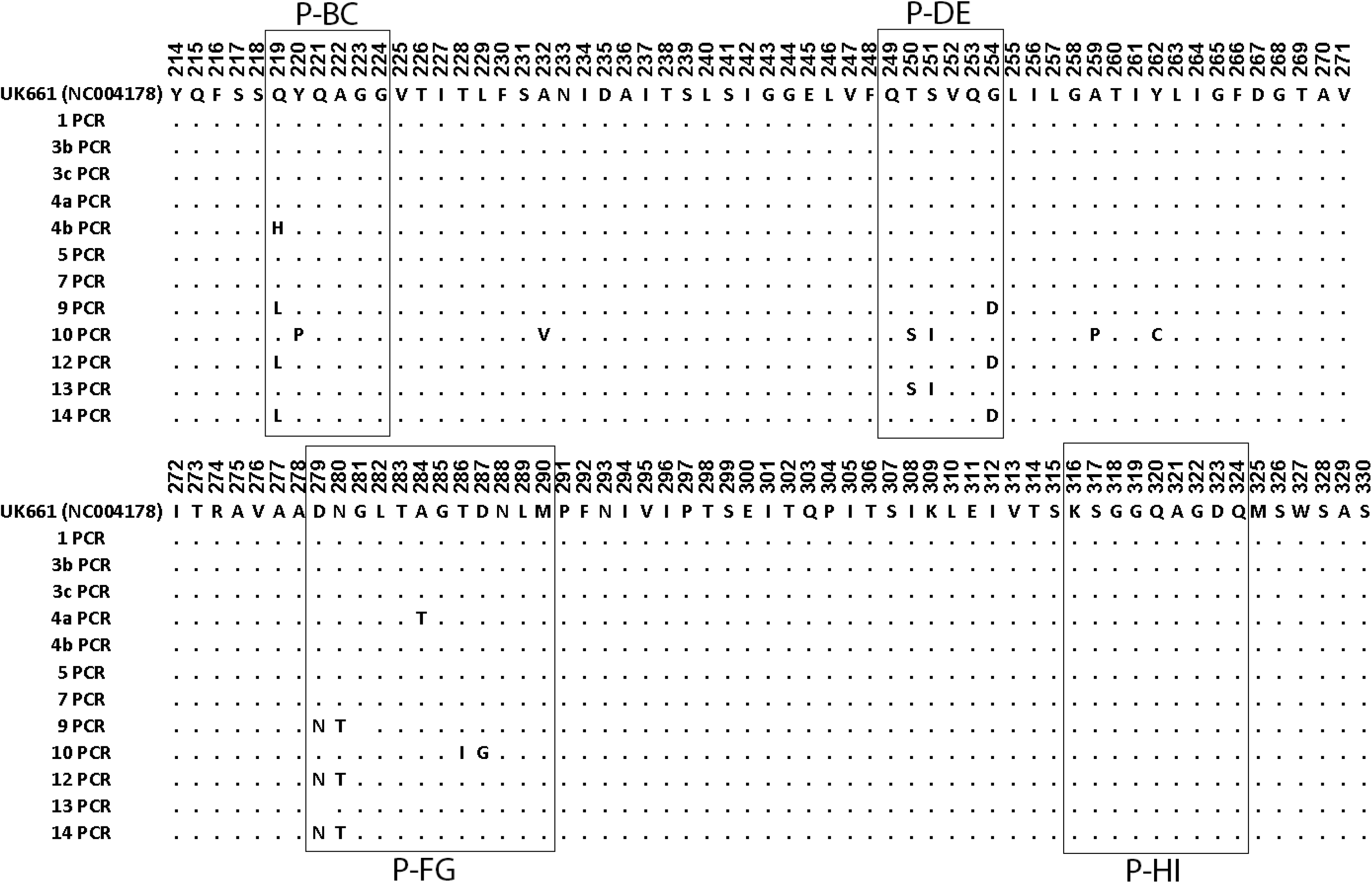
Alignment of the HVR amino acid sequences of the British samples belonging to genogroup A3. The consensus nucleotide sequences of the HVR of the samples that belonged to genogroup A3 were translated *in silico* and the amino acid sequences aligned and compared to strain UK661 (Accession number NC004178). Each amino acid is depicted by its letter abbreviation. Where the amino acid is identical to UK661, it is shown as a dot. The boxes indicate the HVR hydrophilic loops P-BC, P-DE, P-FG, and P-HI.

**Figure 5.**
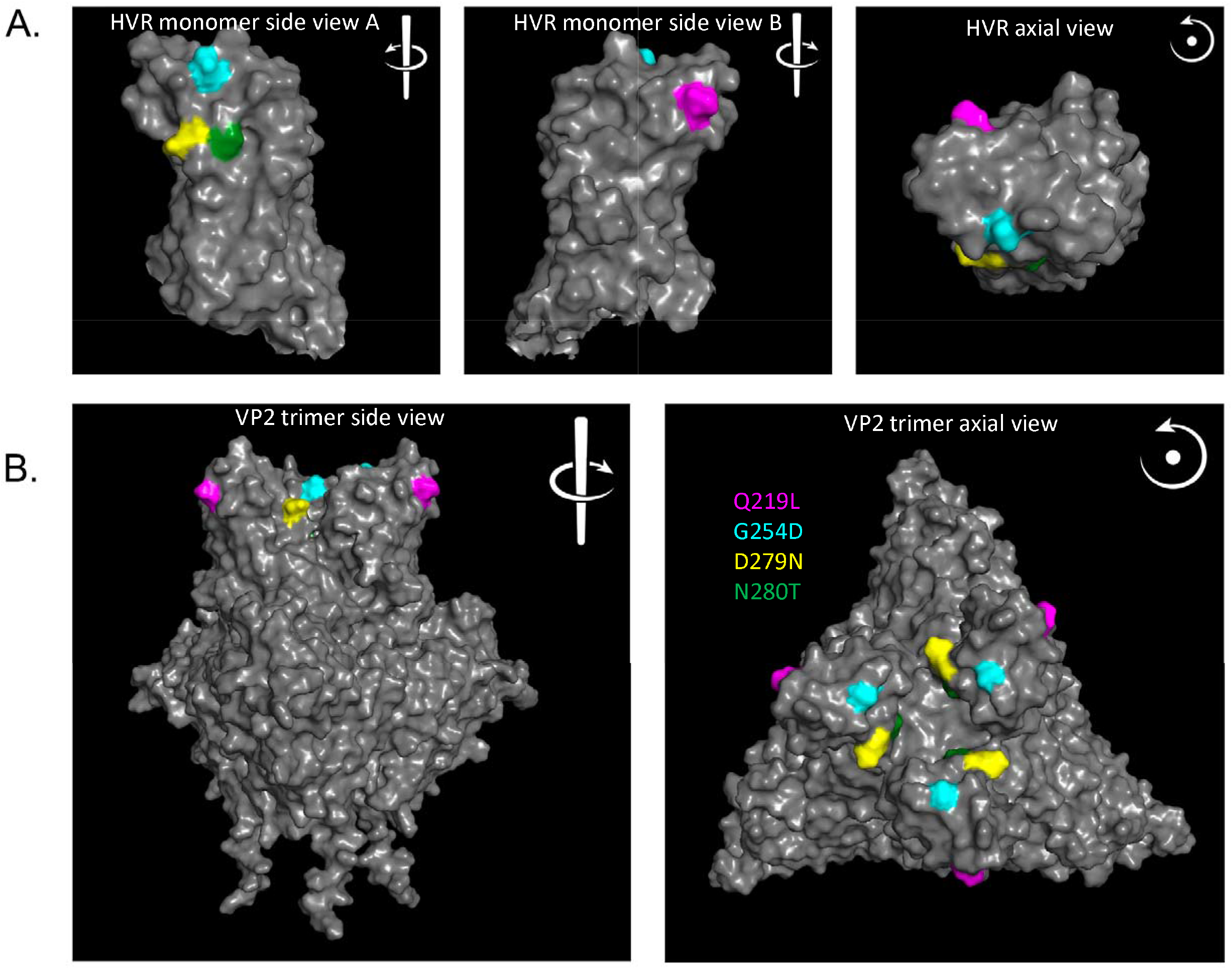
Structural modelling. The predicted structure of the HVR monomer of the Western European clade of reassortant IBDV was modelled using AlphaFold. The predicted structure was loaded and visualized with PyMol, and the side views and end-on / axial view were displayed (A). The VP2 trimer was predicted using the AlphaFold multimer model, and the mutation sites mapped onto the structure (B). The backbone UK661 atoms were depicted as solid grey, and the positions of mutations found in the Western European clade of reassortant viruses were highlighted: Q219L (purple) G254D (blue) D279N (yellow) and N280T (green).

**Figure 6.**
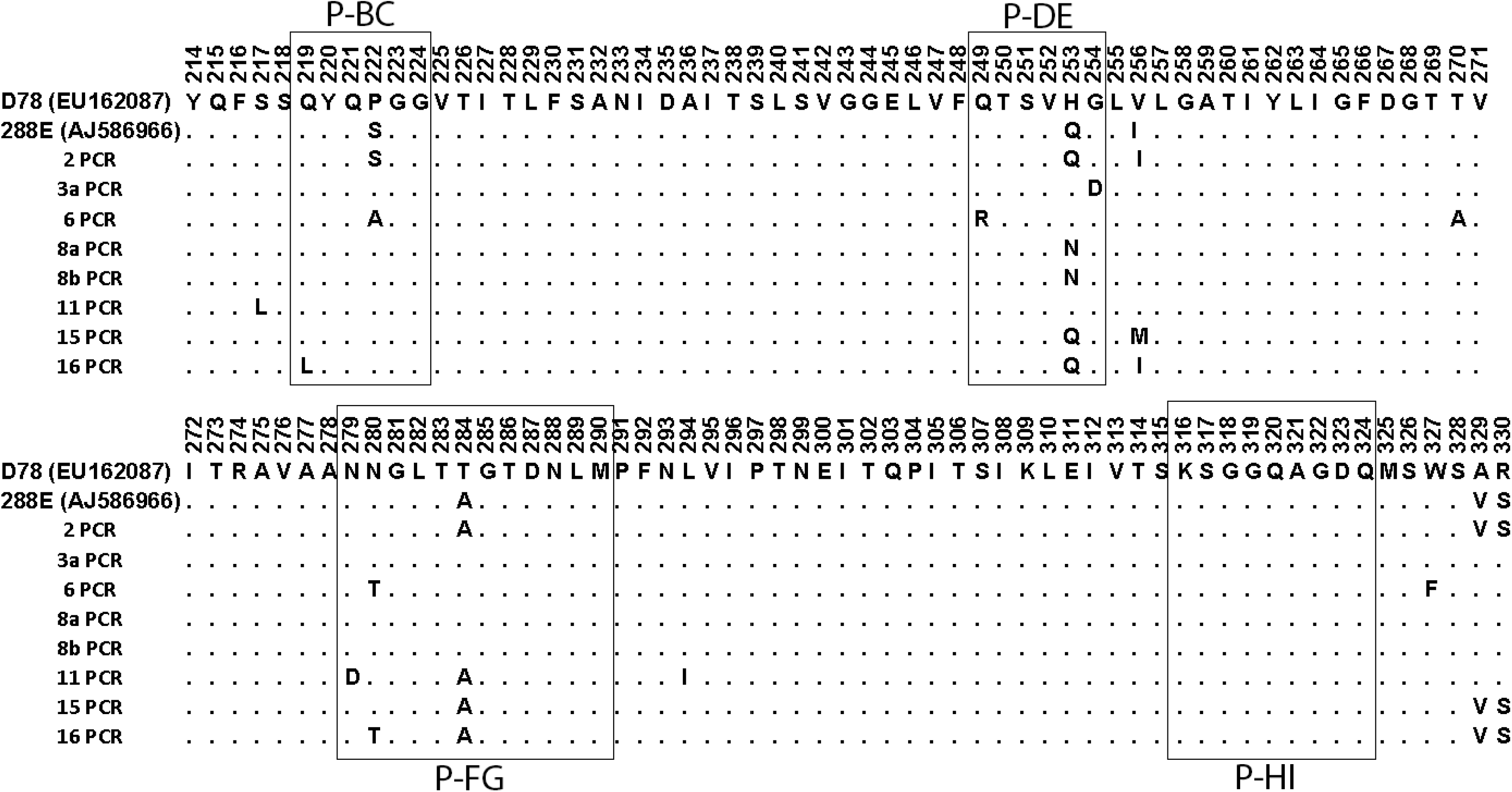
Alignment of the HVR amino acid sequences of the British samples belonging to genogroup A1. The consensus nucleotide sequences of the HVR of the samples that belonged to genogroup A1 were translated *in silico* and the amino acid sequences aligned and compared to strain D78 (Accession number EU162087). Each amino acid is depicted by its letter abbreviation. Where the amino acid is identical to D78, it is shown as a dot. The boxes indicate the HVR hydrophilic loops P-BC, P-DE, P-FG, and P-HI.

**Figure 7.**
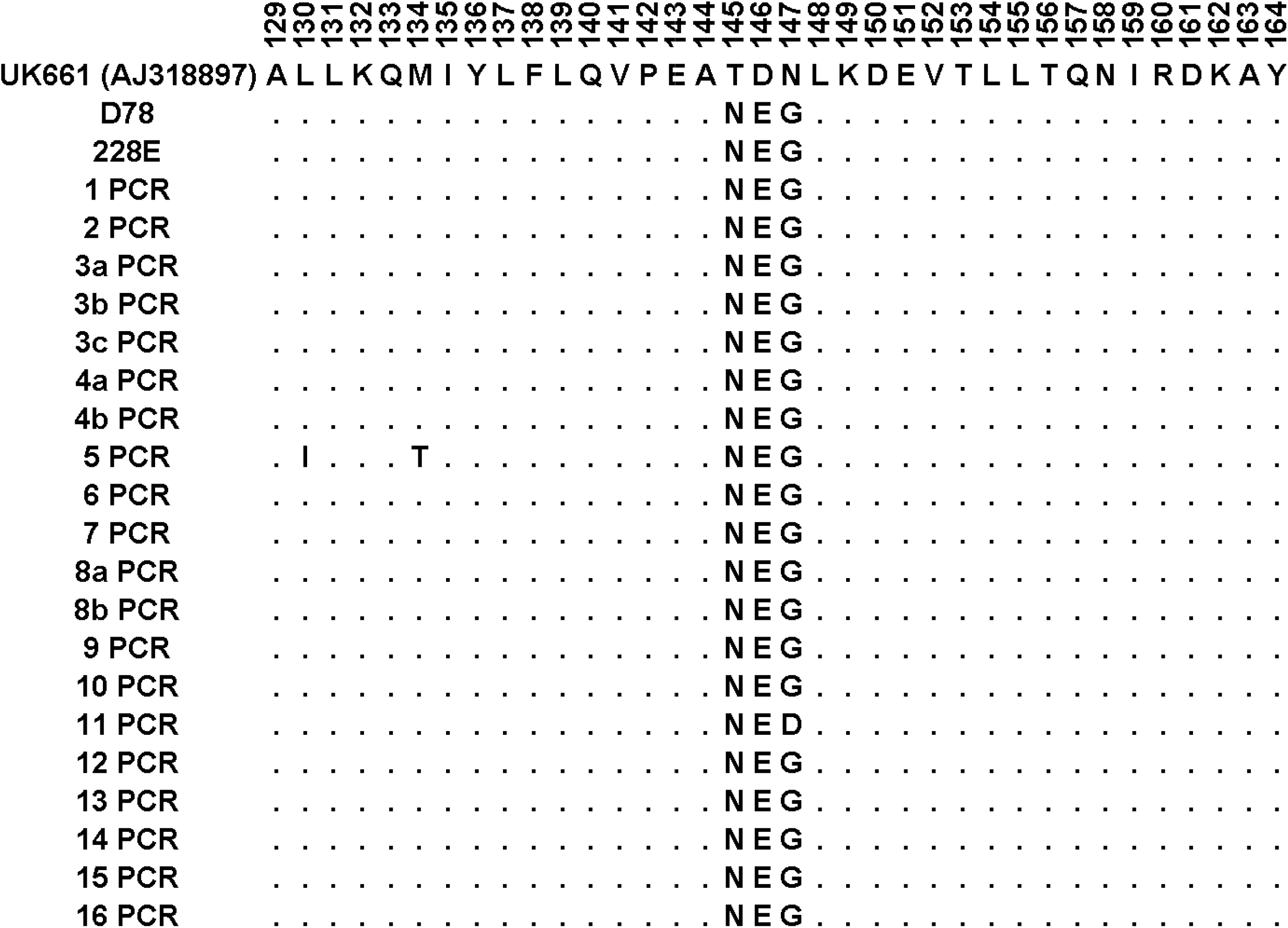
Alignment of the partial VP1 sequences of the British samples. The consensus nucleotide sequences of the VP1 region were translated *in silico* and the amino acid sequences aligned and compared to strain UK661 (Accession number AJ318897.1). Each amino acid is depicted by its letter abbreviation. Where the amino acid is identical to UK661, it is shown as a dot.

## Discussion

As part of ongoing surveillance of endemic poultry diseases in Great Britain, we analysed 20 bursal samples from 16 British broiler farms between 2020 and 2021. During this investigation, we sequenced the VP2 HVR and a region of the VP1 gene from these samples and conducted alignments and constructed phylogenetic trees. While the sample size is limited, the data show that reassortant IBDV strains belonging to genogroup A3B1 were the predominant strain circulating on these farms. IBDV reassortment has been reported with increasing frequency. In 2003, Sun *et al* identified a reassortant strain with a vv segment A and a non-vv segment B in China (25), and in 2006 Le Nouёn *et al* analysed 50 strains from 1972 to 2002 in 17 countries from 4 continents and found that 26% contained incongruency in the phylogenetic trees between segments A and B, suggestive of reassortment, and when one of these was sequenced, it was found to have a vv segment A and non-vv segment B (26). Subsequently, additional reassortant strains with a vv segment and a non-vv segment B have been reported in China (27–30), Brazil (31), Zambia (32), India (33, 34), Algeria (35), Nigeria (36), Europe (9, 16–20), and the Near East and Persian Gulf regions (37). While the majority of these have been genogroup A3B1, there have also been reports of reassortant strains with genogroups A3B3 in Bangladesh (38), A3B4 in Poland (19), and A3B5 in Nigeria (39). In addition, there have been reports of reassortment between IBDV strains with a non-vv segment A and a vv segment B, in China (40, 41), and Egypt (25), and reassortment between other genogroups, for example an A2dB3 strain in China (42). Moreover, there have been two reports of reassortment between strains of different serotype, with a vv segment A and a serotype 2 segment B, in Europe (43), and the US (44).

In Europe, reassortant viruses of genogroup A3B1 were found to be widespread in 2020, present in Latvia, Germany, Belgium, Denmark, The Netherlands, Czech Republic, and Sweden (18). Some of these strains also contained mutations Q219L, G254D, D279N, and N280T in the HVR, demonstrating ongoing evolution by antigenic drift. In 2023, France, Italy, Portugal, Spain, and the UK were added to the countries where the reassortant A3B1 strains were reported, demonstrating that their geographical range was even more extensive than previously thought (16).

In the present study, we extend these observations by demonstrating that reassortant strains belonging to genogroup A3B1 were present in 13/16 (81%) of the British farms sampled. It is therefore possible that these strains are now endemic in British flocks, although a larger sample size is needed to confirm this hypothesis. Moreover, we demonstrate that the majority of farms with reassortant strains were also co-infected with genogroup A1B1 strains, and it is therefore possible that continued reassortment between A1B1 strains and A3B1 strains occurs in the field, although this is difficult to characterize. In addition, based on the partial sequences we obtained, sample 11 contained segment A1 and B1 sequences that were phylogenetically more related to classical field strains, whereas the other segment A1 and B1 sequences were more closely related to vaccine strains. This finding suggests that the reassortant viruses contained B1 sequences derived from vaccine strains. Previously, the Western European reassortant viruses were reported to have a segment B from an attenuated strain (18), and our data are consistent with this report. Moreover, this observation suggests that the A1B1 viruses that co-infected with reassortant strains were also likely to be vaccine strains.

Interestingly, we found no evidence of true vv strains of genogroup A3B2 in our samples, and it is possible that the A3B1 reassortant strains have a fitness advantage over the A3B2 vv strains, thus outcompeting them in the population, although additional studies are needed to confirm this hypothesis. Furthermore, we demonstrated that in 5/13 (38%) of farms with reassortant viruses, the strains had the same genetic signature described in Western European reassortant strains (mutations Q219L, G254D, D279N, and N280T compared to UK661), whereas other reassortant strains had either no mutations compared to UK661, or different point mutations, suggesting that multiple clades of reassortant viruses co-circulate in the UK.

Assessment of the pathogenic potential of the British reassortant viruses was beyond the scope of this study, however, it has previously been demonstrated that the European A3B1 reassortant strains caused subclinical infection with no mortality or clinical signs, but induced a profound bursal atrophy that was presumed to be associated with immunosuppression (18). Consistent with these reports, the flocks from which our samples were obtained did not have an excess in mortality, suggesting the British reassortant strains did not have a very virulent phenotype. The consequences of co-infection of reassortant strains with vaccine or classical strains also remains poorly understood. Given our findings, it will be important to evaluate the pathogenicity and immunosuppressive potential of the reassortant strains alone, and in co-infection studies with genogroup A1B1 viruses, and in vaccinated birds, to simulate the situation emerging in the field. Moreover, it will be important to evaluate differences in pathogenicity and immunosuppressive potential between the different clades of reassortant strains found in the Great Britain. It will also be important to evaluate the extent to which HVR antigenic drift mutations in the reassortant strains contribute to immune escape and vaccine failure. Mutations Q219L, G254D, D279N, and N280T, and mutations T250S and S251I, are present in 3 of the 4 hydrophilic loops on the tip of the HVR (defined as per (11)), which are thought to be the site of most antibody binding (Figure 4). Moreover, previously, we generated IBDV escape mutants *in vitro*, in chicken B cells, and found that one contained both D279N and S251I mutations (45), suggesting that these changes could potentially play a role in escape from immunity. It is, therefore, possible that these mutations contribute to vaccine failure, but this needs to be experimentally tested.

One limitation of our present study is that we relied on sequencing small regions of the VP2 HVR and VP1 only. While this approach has been routinely applied in IBDV epidemiological studies, whole genome sequencing (WGS) of IBDV strains has recently been employed effectively (46, 47). Moreover, the Western European reassortant strains were found to contain mutations elsewhere in the genome, in regions encoding VP3, VP4 and VP5 (18), and it would be beneficial to determine whether these mutations are also present in the British strains.

Finally, during IBDV assembly, the virus particle is nucleated around the viral RNA, instead of the packaging of a pre-assembled genome into a new daughter virion. This phenomenon, in addition to a spacious capsid, leads to some viral particles being polyploid, containing up to 4 genome segments, even though only one segment A and one segment B is needed for the particle to be infectious (48). As we identified segment A3 and A1 genogroups together in 50% of the bursal samples, this could be either co-infection with two distinct viruses, or it could be infection with one virus containing multiple segment As belonging to two different genogroups. Further studies are required to dissect this phenomenon. Never-the-less, the fact that we co-amplified multiple segments in the same sample highlights the need to determine IBDV diversity at the sub-consensus level in epidemiological studies. While this was done in the present study by TA cloning, in the future, next generation sequencing (NGS) technologies should be employed.

In summary, true vv strains of genogroup A3B2 were not detected in the British broiler flocks we sampled, whereas reassortant IBDV strains belonging to genogroup A3B1 predominated. In addition, multiple clades of A3 reassortant viruses were found to be circulating, and co-amplification of A3 and A1 sequences was common, suggesting that reassortant strains were commonly found in birds co-infected with vaccine strains. The consequences of reassortment, antigenic drift, and co-infection in terms of pathogenesis, clinical presentation, immunosuppressive potential, and immune escape remains poorly understood, and further studies are therefore required to improve IBDV disease control.

## Acknowledgements

We acknowledge the farmers and poultry producers who permitted sampling of their flocks, and the private veterinarians and individuals who collected the bursal samples for this study.

## Funding

This work was funded by the Department of Environment, Food and Rural Affairs and The Welsh Government through the “Scanning Surveillance for Disease in Avian Farmed Species in England and Wales” (ED1300) project at APHA, and by projects BBS/E/I/00001845, BBS/E/I00007034, and BBS/E/I/00007039, at The Pirbright Institute, funded by the Biotechnology and Biological Sciences Research Council (BBSRC), UK. The funders had no role in study design, data collection and interpretation, or the decision to submit the work for publication.

## Conflict of Interest

The authors declare no conflict of interest.

## Ethical statement

The considered samples were collected within the context of routine diagnostic activities and not for experimental purposes. No ethical approval was therefore required.

## Notes

### Competing Interest Statement

The authors have declared no competing interest.

